# Enzyme encapsulation peptides bind to the groove between tessellating subunits of the bacterial microcompartment shell

**DOI:** 10.1101/2022.12.30.522206

**Authors:** Jack Bradley-Clarke, Shuang Gu, Ruth-Sarah Rose, Martin J. Warren, Richard W. Pickersgill

## Abstract

Bacterial microcompartments are prokaryotic organelles consisting of encapsulated enzymes within a thin protein shell. They are involved in metabolic processing including propanediol, choline, glycerol and ethanolamine utilization, and carbon fixation. Enzymes targeted to the inside of the microcompartment frequently possess a cargo-encapsulation peptide, but the binding-site has not been revealed. We provide evidence that the encapsulation peptides bind to the hydrophobic groove formed between tessellating tiles of major shell proteins. *In silico* docking studies provide a compelling model of peptide binding to this hydrophobic groove. This result is consistent with the now widely accepted view that the convex side of the shell proteins faces the lumen of the microcompartment. Binding between shell protein tiles explains why the encapsulation peptide binding site has been elusive, how encapsulation-peptide bearing enzymes can promote shell assembly, and how the presence of cargo affects the size and shape of the bacterial microcompartment.

## Introduction

Bacterial microcompartments are prokaryotic organelles consisting of encapsulated enzymes within a thin protein shell. The first bacterial microcompartments, observed as polyhedral structures in electron micrographs, were the carboxysomes of Cyanobacteria^1^ which enhance carbon dioxide fixation via encapsulation of rubisco and carbonic anhydrase^2^. Later, similar structures were observed in heterotrophs, but only when grown on the substrate of the microcompartment e.g., ethanolamine or 1,2-propanediol^3^. The majority of the bacterial microcompartments break down a metabolic substrate and are called metabolosomes. The bacterial microcompartment/metabolosome shell functions as a semipermeable membrane for substrates and products and segregates the encapsulated enzymes^4^. The shell confines toxic and reactive intermediates and enhances catalysis by increasing the concentration of enzymes and substrates. A recent paper catalogues the increasing known diversity and ubiquity of bacterial microcompartments^5^. Targeting of enzyme cargo to the lumen of the bacterial microcompartment is typically by a 15-20 amino-acid residue amphipathic alpha-helix that is connected to the N- or C-termini of the cargo protein via a flexible linker^6,7^.

The propanediol utilization (Pdu) metabolosome from Salmonella comprises eight shell proteins (PduA, B, B’, J, K, N, U, T) of which PduA, B, B’, J are major and PduK, T, U (and N) are minor components of the shell^8^. The shell protein PduA consist of a single Pfam00936 domain that assembles into a cyclic homohexamer with a convex and concave side (Fig. 1)^9,10^. PduB is a tandem fusion of two Pfam00936 domains that assembles into a cyclic homotrimer which closely resembles the size and shape of the PduA hexamer^11–13^. Except for the vertex-capping pentamer, PduN^14^, the shell proteins are either hexamers or pseudo-hexamers. Several thousand of these hexamers and pseudo-hexamers tessellate to form the facets of the bacterial microcompartment.

**Figure 1:**
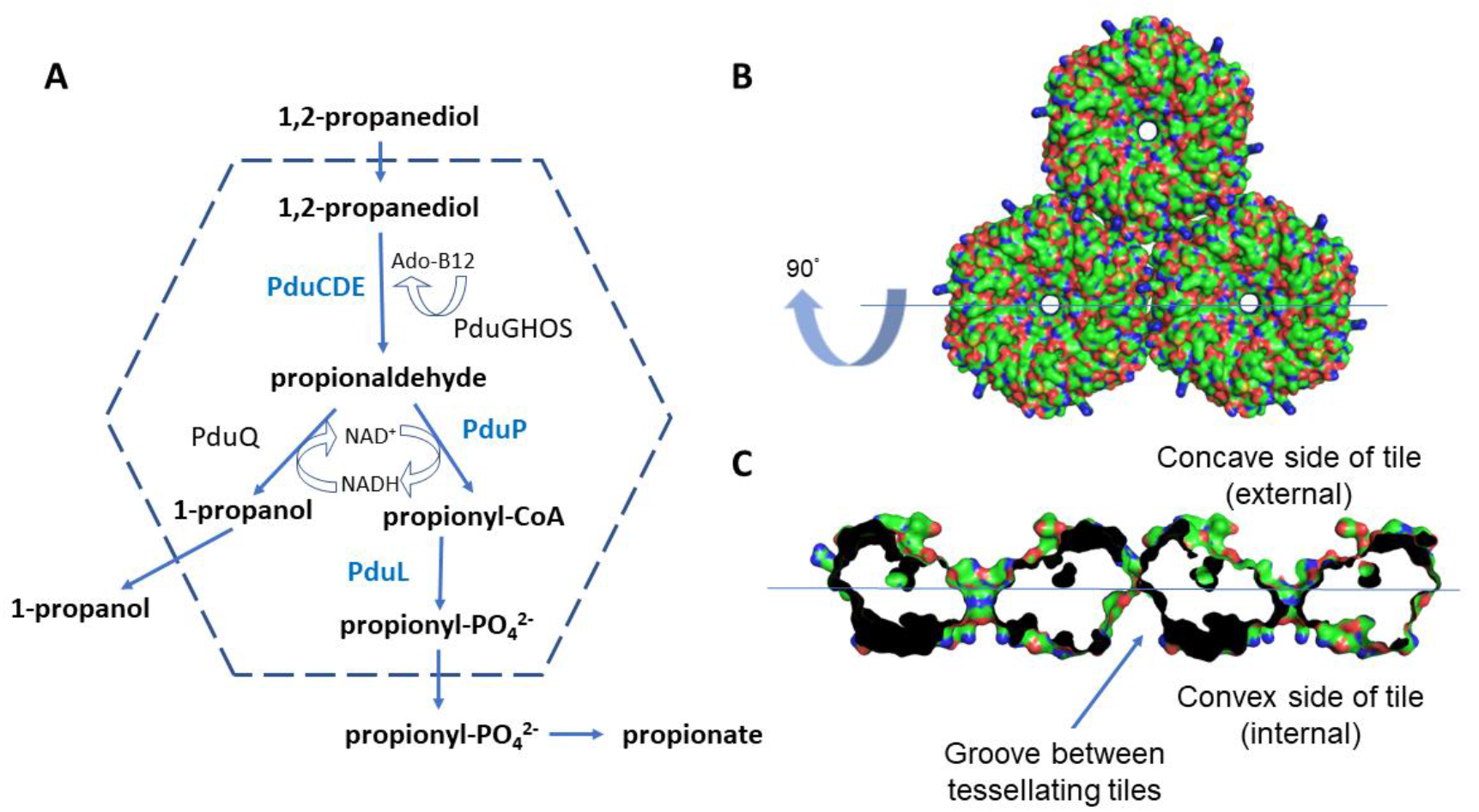
The Pdu microcompartment and tessellating PduA hexamers which form the shell. (A) Schematic representation of the cobalamin dependent 1,2-propanediol utilization (Pdu) microcompartment and the encapsulated enzymes. Enzymes with known encapsulation sequences, PduCDE, PduL and PduP, are highlighted in blue. (B) Three tessellating PduA shell protein hexamers viewed looking down on to the convex face of the hexamers. (C) A central slab of the tessellating PduA molecules rotated about the horizontal axis by 90° compared to panel (B) to show the concave and convex surfaces of the hexamers, the thin hexamer-hexamer interface can be clearly seen between tessellating hexamers. The hydrophobic groove formed between the tessellating PduA protein tiles is indicated on the convex side of the PduA hexamer.

In the Pdu microcompartment, the encapsulated enzymes convert 1,2-propanediol to propionaldehyde via a cobalamin dependent catalytic mechanism catalysed by PduCDE (Fig. 1A). The aldehyde is subsequently converted to propionyl-CoA by PduP^15^ before the CoA is regenerated by PduL during the production of propionyl-phosphate^16^. There is evidence that the aldehyde dehydrogenase (PduP) and the diol dehydratase (PduCDE) are within the lumen of the microcompartment^17,18^. The requirement for PduL to regenerate CoA for PduP would imply that it too is localised to the microcompartment lumen^19^. Several other enzymes (PduGHOS and PduQ) are involved in regenerating the cobalamin and NAD^+^ cofactors. Recombinant production of bacterial microcompartments has shown that in the presence of encapsulation peptide bearing metabolic enzymes the microcompartments are larger compared to empty shells^18^.

In this work we use the hexameric shell protein PduA from *Salmonella typhimurium* LT2. PduA has been shown to transport the substrate 1,2-propanediol to the lumen of the Pdu microcompartment^94^. While PduA comprises only 19% of the shell proteins present in the Pdu microcompartment shell its sequence is closely homologous to PduJ (80% sequence identity) which accounts for 54% of the shell proteins^20^. The structure of PduJ has been shown to be nearly identical to PduA and the *pduA* gene complements the growth phenotype of a *pduJ* deletion mutant^21^. Interestingly, it is the genomic position of the *pduJ* gene in the operon that determines its ability to act as a pore for 1,2-propanediol transport as the substrate channel of PduJ is identical to that of PduA. Together PduA and PduJ account for 73% of the microcompartment facet and we therefore argue that the interactions we establish here using PduA are applicable to the facet of the microcompartment.

Enzymes encapsulated within the Pdu microcompartment have a short, typically 20 residue, encapsulation sequence^22^. These sequences form amphipathic helices with small hydrophobic residues clustered on one side of the helix; they are found at the N-terminus or C-terminus of the enzyme-cargo^23^. Within the Pdu metabolosome, the acylating propanol dehydrogenase, PduP has an 18 residue N-terminal sequence which facilitates encapsulation. Several computational methods have predicted that the encapsulation peptide of PduP binds to the concave surface of PduA, thus requiring the concave side of the PduA tile to be luminal^24,25^. However, there is now convincing evidence that the concave surface of the PduA tile is external both from our own work^26^ and from structural studies of a recombinantly generated metabolosome^27^. While there may be some affinity for the concave surface of the tile, the work we report here reveals a greater affinity for the hydrophobic groove formed between tessellating tiles. Enzymes bearing encapsulation peptides are not essential for microcompartment assembly^28,18^ but they do influence assembly. The binding of the encapsulation peptide between tessellating tiles has important consequences for understanding the nucleation of bacterial microcompartment assembly in the presence of cargo-enzymes and provides an explanation of how enzyme-cargo influences microcompartment size and shape.

## Results

### PduA tessellation intermediates

PduA hexamers tessellate to produce protein sheets and nanotubes^10,26^. Substitution of residues at the tessellation interface, including the key residues lysine 26 and arginine 79, reduce the propensity of PduA to tesselate^10^. The tessellation of PduA hexamers is also pH dependent and, here, we use low pH and sonication to disrupt the sheets and nanotubes and produce assembly intermediates. A glutaraldehyde cross-linked sample of sonicated PduA was analysed by size-exclusion chromatography (Fig. 2A) and can be seen to comprise three distinct peaks which are interpreted to be a trimer of hexamers, a dimer of hexamers and the single hexamer (Fig. 2B). The peaks fractions are predominantly trimer and dimer (Peak 1) and monomer of hexamers (Peak 2). Dynamic light scattering of samples from the peaks have hydrodynamic radii of 6.68nm and 3.71nm consistent with the measured maximal radius of a tessellated cyclic PduA trimer of hexamers or a dimer (Peak 1), or a single hexamer (Peak 2) respectively (Fig. 2A). The size from dynamic light scattering is confirmed by mass-spectrometry where the masses of 88kDa, 176KDa, 264kDa (Fig. 2C) which are consistent with these assembly intermediates being dimer (176kDa) and trimer of hexamers (246kDa). On SDS-PAGE the glutaraldehyde modified PduA runs anomalously because the sample is not thoroughly unfolded due to the glutaraldehyde cross-links. PduA bands can be seen at approximately 60kDa, 120kDa and 190kDa which are smaller than those measured by mass-spectrometry, but which do support a ladder of oligomers (Fig. 2D). The compact trimer of dimers (as shown in the Figure) is preferred to a linear trimer as the latter would be less stable with only two interfaces compared to the three hexamer-hexamer interfaces in the cyclic trimer. PduA without cross-linking (Fig. 2B) tends to assemble into a cyclic trimer of hexamers.

**Figure 2.**
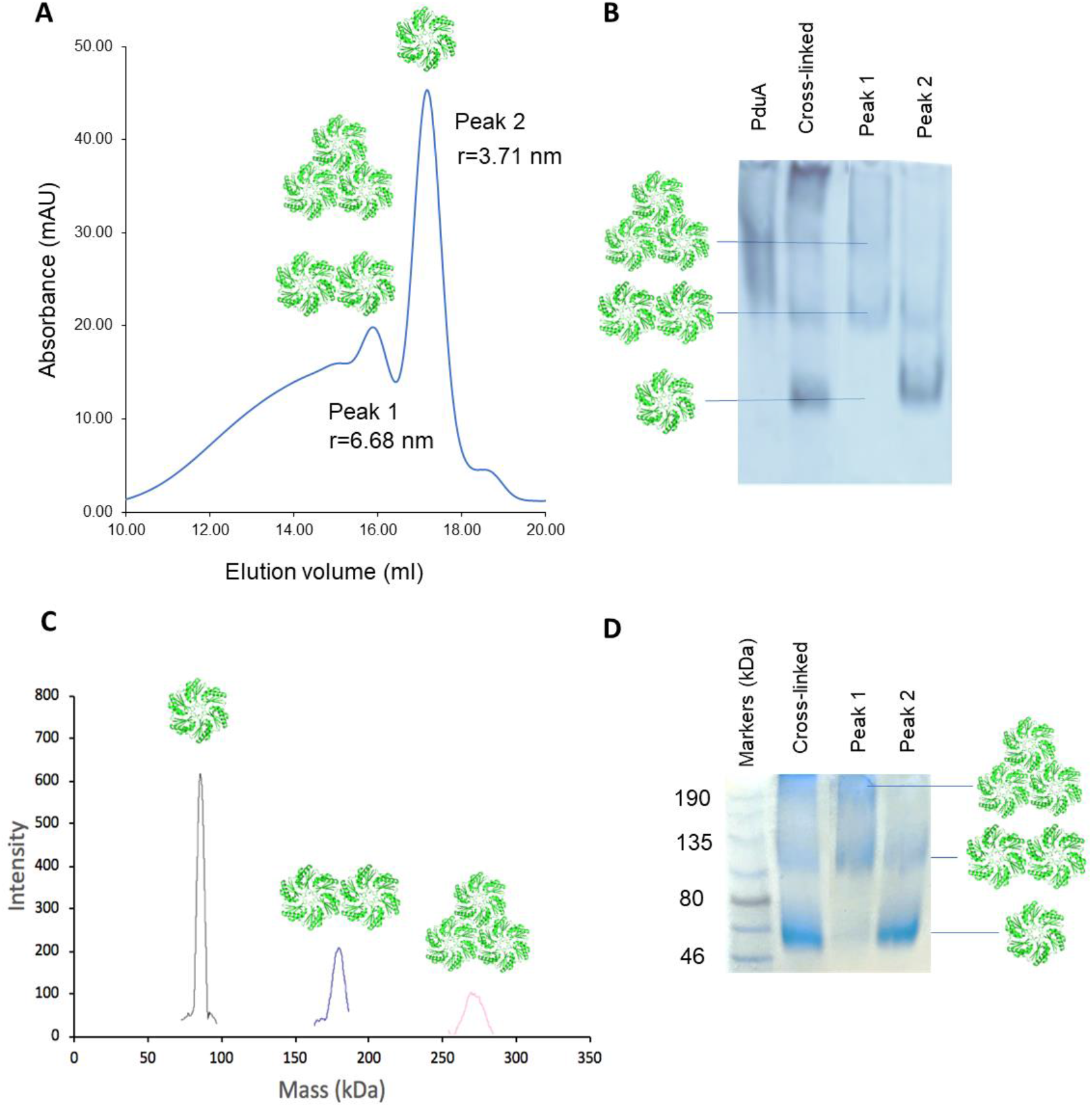
Identifying early-stage tessellation intermediates. (A) After sonication and cross-linking, low molecular mass oligomers of PduA hexamers can be separated using size-exclusion chromatography. Dynamic light scattering measurements suggest that the first peak corresponds to dimers and trimers of hexamers (r=6.68nm) while the second peak is predominantly the PduA hexamer (r=3.71nm). (B) The native gel is consistent with peak 2 comprising mostly monomer, peak 1 trimer and dimer. The glutaraldehyde cross-linked sample contains all three components, monomer, dimer, and trimer. The PduA samples used for the work described here were checked and were predominantly trimers of hexamers (first lane of gel). (C) The mass of the peaks from the glutaraldehyde cross-linked sample was confirmed by MALDI-TOF mass-spectrometry. (D) Cross-linked samples run on SDS-PAGE gives molecular masses in agreement with the results of dynamic light scattering and mass-spectrometry.

### PduL binds to tessellating PduA

We explored the binding of PduL to tessellating and mutated non-tessellating variants of PduA. A distinct band shift is observed on the native gel when PduL is titrated into tessellating PduA hexamers but not when added to the non-tessellating mutant, K26D PduA (Fig. 3A). As expected, the non-tessellating K26D PduA has higher mobility than the tessellating native PduA because of its lower mass and greater negative charge. The increase in the mobility of PduA on addition of PduL was not expected. It is unlikely that the addition of PduL is breaking up the stable PduA hexamer into a lower mass species. More plausible is that the negative charge of PduL (pI 6.2) is pulling the complexed PduA (pI 8.0) further into the gel which has a running pH of 8.0. The non-tessellating variant 6A was also used to confirm the result (Fig. 3B). In this variant the six chains of the hexamer are concatenated into a single polypeptide chain with linkers between the six concatenated subunits. When PduL is titrated into A6 no interaction is seen with PduL (Fig. 3B). This shows that PduL does not interact with the concatenated form presumably because the linker used to make A6 interferes with the PduL binding-site. These variants do not tessellate so this result again links tessellation and PduL-binding. The behaviour of the slowly tessellating PduA A6 variant, (K26D)4, is interesting (Fig. 3C). This variant has four of the six concatenated PduA copies with aspartate in place of lysine 26 and two retain the original lysine. What can be seen is that on addition of PduL the tessellating fraction of the sample undergoes a band-shift, but the non-tessellating component of the sample does not (Fig. 3C). This result strongly supports the view that tessellation is needed for PduL binding. A titration of PduL into PduA reveals a 1:1 binding stoichiometry (Fig. 3D upper panel). The Western blot (Fig. 3D lower panel) shows PduA is present in the lower band on the native gel, revealing that this band is the complex of PduL and PduA.

**Figure 3:**
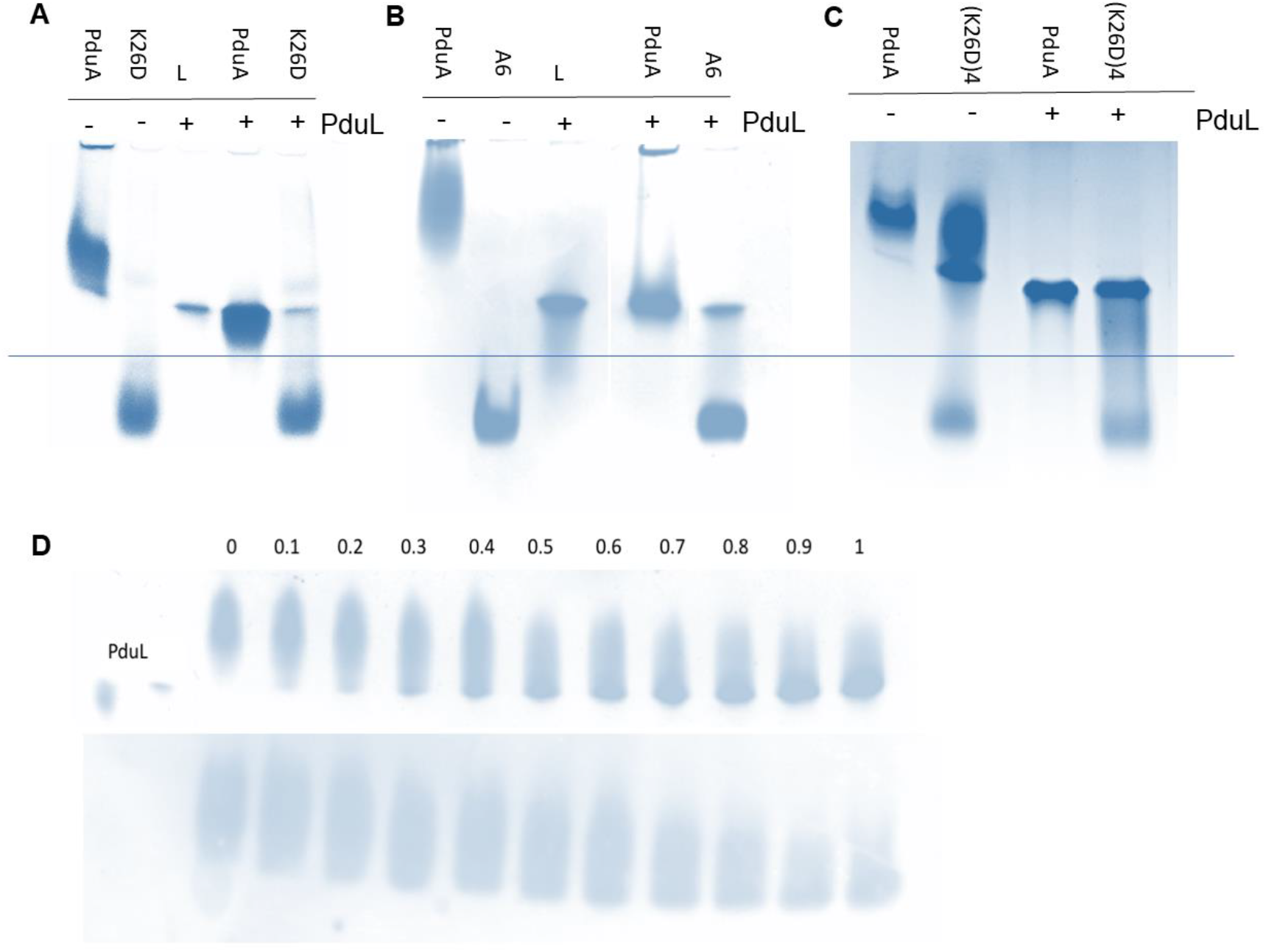
PduL binds only to tessellating PduA hexamers. Tessellating PduA has lower mobility on native PAGE and is observed above the horizontal blue line on the figure and the band shifts when PduL is added, non-tessellating PduA has higher mobility and is seen below the blue line. (A) PduA and PduA variants without and with the addition of PduL. The non-tessellating variant, K26D with the interfacial lysine 26 replaced by aspartate, has lower mass and correspondingly higher mobility. Addition of the more acidic PduL has a profound influence on the mobility of the PduA trimer, but not on that of the K26D mutant, where both PduA and PduL bands are unchanged. (B) A similar result to that presented in the first panel is seen when the concatenated 6A PduA variant is used. This PduA variant does not tessellate and comprises six fused subunits in a single polypeptide chain, the individual subunits joined by linkers. Two distinct bands corresponding to non-interacting PduA variant A6 and PduL are seen indicating no binding. (C) The (K26D)_4_ mutant of PduA, in which four of the six lysine 26’s are replaced by aspartate, tessellates slowly (over several days). Here is a sample that has been left for two days (lane 2). Some of the protein has tessellated (low mobility), some is still non-tessellating (high mobility). What is striking here is that the tessellating species is binds PduL and is pulled further into the gel, while the non-tessellating species remains unchanged. Native PduA behaves as usual and shows the usual band shift on addition of PduL. (D) Titration of PduL into PduA confirms a stoichiometric binding ratio. Native gel (upper) and Western blot (lower) stained using an antibody to PduA.

### *In silico* modelling of encapsulation peptide binding

The binding of the cargo encapsulation peptides of PduL (L20), PduD (D18) and PduP (P18) to the PduA hexamer and dimer of hexamers was evaluated *in silico* using several docking methods: ClusPro^29^, Frodock^30^ and CABS-dock server^31^. The peptides were modelled both as helical and as flexible peptides and the search extensively covered the surface of the hexamer and the dimer of hexamers. When the surface of the monomer is searched, the peptides bind to the concave side of the PduA disk in the mode described previously^22^ (Fig. 4A). The second ranked hit is substantially the same as the top hit, but the third-ranked it is on the convex-side close to the hexamer-hexamer interface (Fig. 4B). When the surface of tessellating PduA hexamers is searched, the results consistently showed binding to the groove at the hexamer-hexamer interface rather than the surface of the hexamer (Fig. 4C). For instance, the rmsd for L20 binding to the hexamer-hexamer interface, using CABS-dock, was 0.88 Å with a cluster density of 113 (compared to the significantly poorer values of 3.0 and 27 for binding to a single hexamer). In this CABS-DOCK model, the irregular starting structure is predicted to bind to the groove as an amphipathic helix (Fig. 4C). In summary, the encapsulation peptides are predicted to bind strongly to the groove between tessellating hexamers on the convex side of the hexamers (Fig. 4C, D).

**Figure 4:**
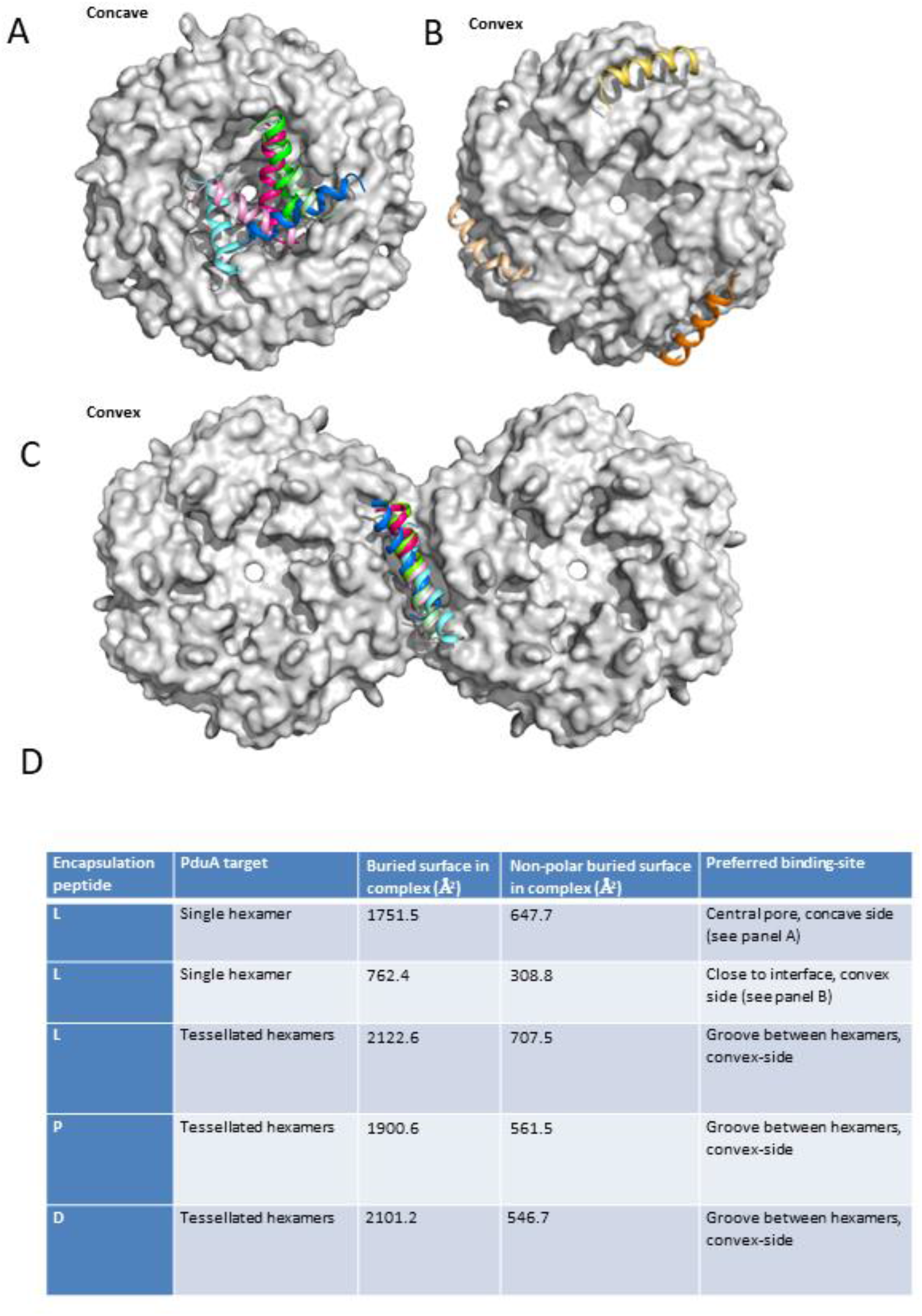
*In silico* modelling of the structure of the encapsulating peptide binding to the PduA hexamer and to tessellating hexamers. (A) Top two hits from docking the encapsulation peptides with the PduA hexamer: L20 (the first 20 residues of PduL in light and dark pink), P18 (light and dark green) and D18 (light and dark blue). The binding is to the concave side of PduA. (B) The third ranked hits for all three encapsulating peptides bind to the convex side of the hexamer close to the hexamer-hexamer-interface. (C) When presented with tessellated hexamers, despite given no preferred binding site residues on PduA, all three encapsulation peptides localise to the cleft between hexamers on the convex side of the PduA dimer. (D) A summary of peptide binding to PduA hexamer and tessellating hexamers.

### Titration of the PduL dimer into the PduA trimer of hexamers

The stoichiometry of binding of 1:1 for the PduL dimer to PduA trimer of hexamers was established using the 6% acrylamide gel (Fig. 3D). The binding was explored at higher resolution using a gradient native gel. The lower mobility band on this gel corresponds to PduA trimer of hexamers and the highest mobility band to the stoichiometric complex of three PduL dimers per PduA trimer of hexamers (Fig. 5A). As the ratio of PduL increases across the gel from left to right, two bands are observed between the empty and fully saturated PduA trimer of hexamers. We interpret the intermediate bands as the binding of one and two PduL dimers per PduA trimer of hexamers (Fig. 5A). In contrast to the nanotubes seen in electron microscopy when using PduA alone, in the presence of PduL or other encapsulation peptides only sheets are seen (supplementary Fig. S1). This is consistent with the cargo-encapsulation peptide binding between hexamers because when the binding-site is occupied by peptide the hexamer-hexamer interface cannot bend to the angle required to make a tube^26^. The apparent cooperativity of binding is interesting (Fig. 5A), and is considered in the discussion, it is consistent with binding at or close to the hexamer-hexamer interface.

**Figure 5.**
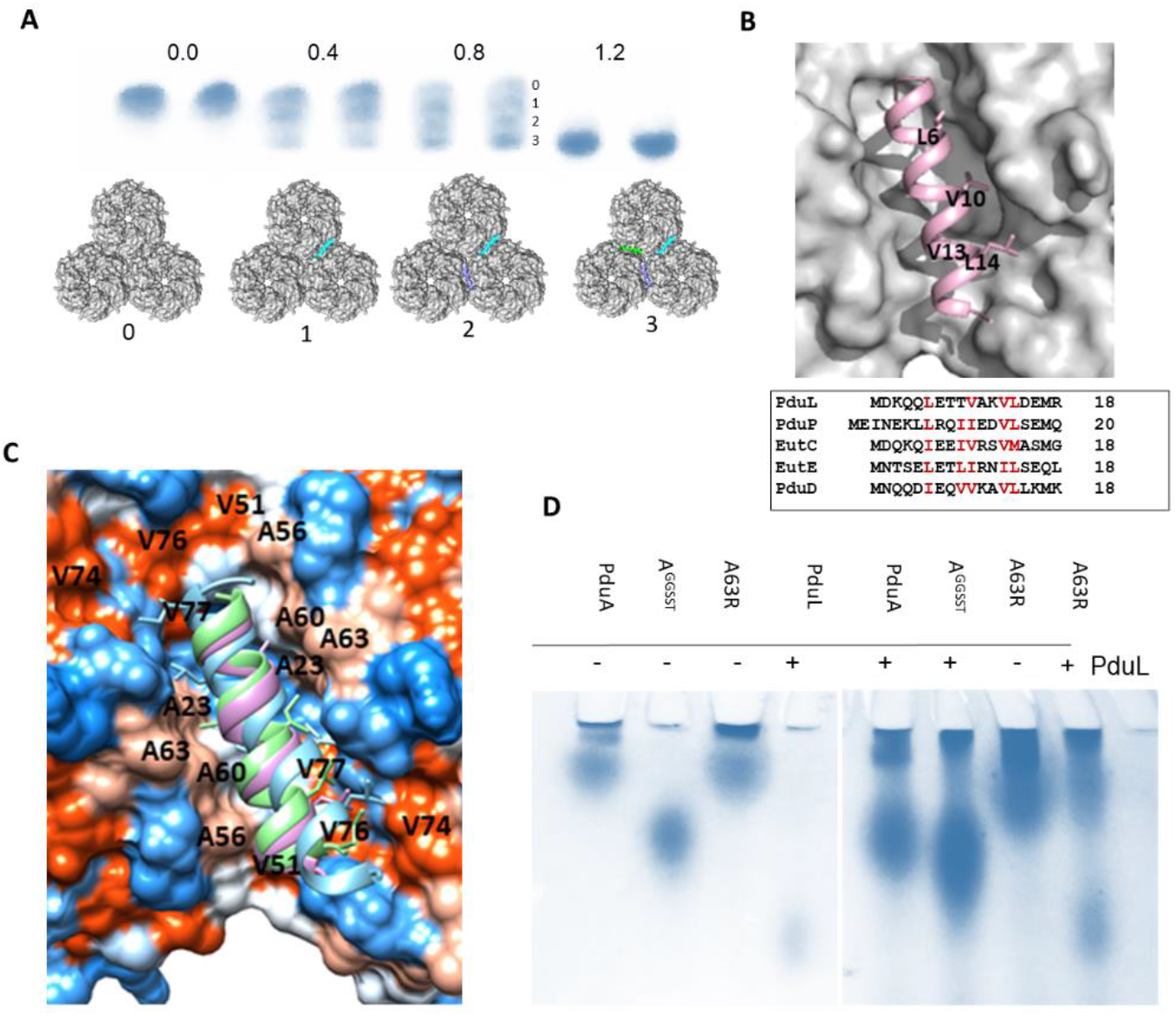
Further exploration of PduL binding to PduA. (A) The top row gives the ratio of PduL to PduA below which is the experimental result. This is a gradient native gel showing the shift from PduA hexamers (left pair of columns; 0.0 PduA) to a stoichiometric ratio of PduL to PduA (right pair of columns). The intermediate two pairs of columns show the progress during titration with four species, labelled 0 to 3, clearly seen in the third pair of columns of the panel. These bands correspond to PduA trimer with none (labelled 0), with one (labelled 1), with two (labelled 2) and with three PduL (labelled 3) dimers bound as illustrated bound to the surface of the PduA trimer of hexamers on the bottom row of the panel. (B) Closeup of the encapsulation peptide binding site between tessellating hexamers of PduA. The conserved hydrophobics which form a patch on the surface of the helical encapsulation peptides are shown as sticks and labelled. (C) The position of conserved small hydrophobics on the surface of PduA on α-helix 1 (<u>A</u>MVK<u>S</u>A; residues 23 and 27 underlined) and α-helix 2 (<u>A</u>ATD<u>A</u>GA<u>A</u>AA; 56, 60 and 63 underlined). The mutation A63R was designed to block the binding site. (D) The result of the band-shift assay using the PduA A63R mutant. Both PduA and A63R can be seen to tessellate (running high on the gel). The PduA A63R mutant is however unable to bind PduL whereas PduA undergoes the usual band shift. PduA with C-terminal linker, GGSST, is used here as a non-tessellating control, it does not bind PduA.

### A mutation that allows PduA tessellation but prevents PduL binding

The binding of the encapsulation peptide to the tessellation interface is shown in Fig. 5B. Three mutations were made to introduce bulky groups to block the PduL binding site identified in the modelling studies (Fig. 5C). One of these three mutants, A63R, was seen to form sheets using electron microscopy so tessellation of the PduA hexamers is preserved. This mutant is more basic than PduA so required the native gel to be run at pH 9.5 instead of 8.5. Compared to the non-tessellating mutant with the C-terminal extension (GGSST), A63R has identical mobility to PduA confirming that it can successfully tessellate to form a trimer of hexamers (Fig. 5D). When PduL is added to A63R no binding is observed with the two bands, PduL and A63R, running as separate bands with unchanged mobility. This contrasts with the band shift seen with native PduA (Fig. 5D). This result reveals that if the groove formed between tessellating hexamers is blocked then it is unable to bind the PduL encapsulation peptide.

### Conformational change in the PduL encapsulation peptide (L20) on binding PduA

To investigate the structure of the PduL encapsulation peptide, the protein from *Citrobacter freundii* was crystallised and its structure determined to a resolution of 1.84 Å. However, the N-terminal encapsulating residues were not seen in the electron density map, a result consistent with the encapsulation peptide being unstructured and flexible. Circular dichroism spectroscopy reveals the P20 and D18 to be helical in solution. The L20 peptide is unstructured in solution though its helicity increased on binding to PduA but did not increase when added to non-tessellating PduA variant (Supplementary Fig. S2). P18 has previously been shown to have helical propensity^32^.

## Discussion

Using gentle sonication to disrupt higher order assemblies and higher pH to slow subsequent reassembly we have been able to separate early-assembly intermediates in the formation of PduA sheets and nanotubes. Frequently observed, the higher molecular mass assembly of PduA hexamers travels more slowly through the native gel than variants that do not tesselate. Tessellation of PduA was confirmed using electron microscopy and observing the presence of sheets and nanotubes. These higher-order structures were not seen using non-tessellating variants, and nanotubes were not seen when encapsulation peptides were added, and the bending of the hexamer-hexamer interface thereby restricted. The planar cyclic trimer with 3-fold symmetry is proposed to be the favoured assembly intermediate because it is the most compact arrangement where each of the three tiles is stabilised by the interaction with two adjacent tiles. The non-tessellating variants of PduA used in this study have mutations near the hexamer-hexamer interface or the addition of flexible linkers that are presumed to interfere with the hexamer-hexamer interface. Unlike tessellating PduA, these non-tessellating variants do not bind PduL. This is either because the cargo-encapsulation peptide binds close to the hexamer-hexamer interface or because the encapsulation peptide binds in the hydrophobic groove formed between adjacent tessellating hexamers. The correlation of tessellation and cargo-encapsulation peptide binding is striking and established using several mutants of PduA including the slowly tessellating mutant where only the tessellated oligomer binds to PduL. These experiments provide a clear and direct link between PduA tessellation and PduL binding.

*In silico* studies suggest the binding of the cargo-encapsulation peptides is preferentially to the hydrophobic grooves between the tessellating hexamers. The peptides adopt helical conformation on binding and present a hydrophobic surface to the binding site. This mode of binding would not be readily detected in previous *in silico* studies using isolated hexamers possessing half the binding site. The conserved hydrophobic residues of the helical encapsulation peptides interact with small hydrophobic residues on the α1 and α2-helices of two adjacent PduA hexamers. The residues on the first helix are: 23 and 27; and on the second: 56, 60 and 63. Mutation of one of these hydrophobic residues, Ala 63, to the bulkier and charged arginine blocks the binding cleft, transforms its hydrophobic character, and prevents the binding of the encapsulation peptide despite tessellation of the PduA hexamers. This mutagenesis result supports the *in-silico* modelling and binding of cargo-peptide to the hydrophobic groove between tessellating tiles. Binding of the encapsulation peptide is to the convex side of the hexameric disk in agreement with previous studies showing this side of the hexamer faces the lumen of the microcompartment.

Further evidence for binding between tessellating hexamers comes from the higher-resolution titration which shows saturation at three PduL dimers per PduA trimer consistent with the binding to the three hexamer-hexamer interfaces present in the trimer. A plausible explanation for the positive cooperativity seen is that the binding of two encapsulation peptides flattens the assembly of three hexamers, opens the third binding groove, and thereby increases the affinity for the third peptide. Electron microscopy shows that the presence of the encapsulation peptides inhibits the formation of PduA nanotubes which require bending at the junction between hexamers, but as expected the presence of encapsulation peptide does not prevent the formation of flat protein sheets. We can now understand how the encapsulation peptide influences the assembly of the microcompartment. Binding to a single hexamer would not directly affect assembly, except by influencing the effective concentration in the vicinity of the enzyme, but binding between hexamers will stabilise the hexamer-hexamer interface, increase facet stability and planarity and directly promote microcompartment assembly. This binding may contribute to determining which tiles are close to a particular encapsulated enzyme. The groove between tessellating hexamers has two-fold symmetry, this greatly complicates achieving ordered binding of peptides for experimental structural studies when using arrays of tiles. When the encapsulation peptides are attached to their cargo enzymes, the oligomeric state of the cargo and the steric exclusion of the enzymes will affect how the peptides are presented and it is plausible this will influence binding to higher order arrays.

A groove between tessellating tiles is a common theme in the crystal structures of shell proteins and the sequence of small hydrophobics of α-helix 1 (<u>A</u>MVK<u>A</u>A; residues 23 and 27 underlined) and α-helix 2 (<u>A</u>ATD<u>A</u>GA<u>A</u>AA; 56, 60 and 63 underlined, 3NGK numbering) that form the binding-site is conserved across many hexameric shell proteins including the major shell proteins PduJ and PduA (Supplementary Fig. S3). This hydrophobic groove is therefore present in the facets of the bacterial microcompartment shell. We suggest that encapsulation peptide binding to grooves formed between tessellating subunits is a general way of binding cargo and plausibly also of prompting nucleation of the microcompartment shell.

## Methods

### Molecular biology

We previously used a variant of PduA with a C-terminal 23 residue extension which aids protein solubility. This is the variant used here too, it is referred to in the methods as PduA*, but simply as PduA in the main text. The constructs PduA*, PduA*_K26D_ and PduA*_R79A_ in pET14b have been described previously^10^. The concatenated constructs of 6xPduA*_4R79A,2WT_ and 6xPduA*_4K26D,2WT_ were prepared in pOPIN F (OPPF) modified to contain a TEV cleavage site and the restriction sites SpeI, EcoRV and BglII (gift) inserted into the KpnI site. PduA* or PduA*_K26D_ were amplified using primers containing ScaI in the forward primer (5’-3’) gg<u>agtact</u>atgcaacaagaagcgttagg and incorporating EcoRV and BglII sites either side of a stop codon using the primers (5’-3’) at<u>agatct</u>tta<u>gatatc</u>ttgctcagcggtggcagc. The PCR product was ligated into pBluescript SKII +. The gene was excised using ScaI and BglII and was ligated into pOPIN F TEV linearised with EcoRV and BglII. This was repeated in a link and lock style ^33^ approach until 6 copies of PduA, with the * tag between each repeated, had been ligated. The sequence was confirmed after each step (Source BioScience). PduA_GGSST_ was designed to include a 30 residue C-terminal extension, in place of the *tag and was synthesised including the same ScaI, EcoRV and BglII sites as above. The synthesised gene was excised using ScaI and BglII and ligated into pOPIN F TEV linearised with EcoRV and BglII. PduA*_A63R_ was created using site directed mutagenesis.

### Protein production and purification

For protein production, BL21*DE3) transformed with the desired plasmid were grown in 1 litre volumes of 2YT media supplemented with ampicillin, at 37°C while shaking at 200rpm. Gene expression was induced at an OD of 1.0 with 0.4mM IPTG followed by overnight incubation at 18°C, shaking at 200rpm. Cells were harvested by centrifugation at 6000*xg* for 10 mins and were resuspended in 20mM Tris pH 8.0, 500mM NaCl. The cells were lysed by sonication and the lysate clarified using centrifugation at 25000*xg* for 30 mins. Proteins were purified using immobilised nickel affinity chromatography. PduA and mutants were washed with 20mM Tris pH 8.0, 500mM NaCl and imidazole up to 150mM before elution with 500mM. PduL was washed with 20mM Tris pH 8.0, 500mM NaCl and imidazole up to 60mM before elution with 250mM. The N-terminal His-tag of PduL was cleaved after incubation with thrombin at 4°C overnight. Thrombin and un-cleaved protein were removed using reverse immobilised nickel affinity chromatography. The proteins were further purified using size exclusion chromatography on a Superdex 200 10/300 column equilibrated in 25mM HEPES, 500mM NaCl, pH8.0 and eluted at their expected sizes.

### Native-PAGE analysis

For all Native-PAGE, BioRad Mini-PROTEAN TGX 4-15% gradient gels and running buffer (25mM TRIS, 192mM glycine) were run at 4°C for 3 hours at 100V fixed with variable current. Native-PAGE samples were prepared in 200mM NaCl, 25mM HEPES, pH8.0 and protein complexes were left at 4°C for 1 hour before the addition of loading buffer (0.1% bromophenol blue, 50% glycerol, 50% 1x running buffer).

### Cross-linking

To prepare PduA* oligomers, 4 mg/ml PduA* in 20mM HEPES, 500mM NaCl, pH8, was sonicated, on ice, for 30 sec pulses for 2 mins. Glutaraldehyde to 1 % v/v was added immediately after sonication and the crosslinking reaction was incubated 4°C overnight. The reaction was terminated using SEC with Superose 6 column in 20mM Tris, 500mM NaCl pH8.

### Circular Dichroism

PduA* and L20 were prepared in 25mM PO_4_^2-^ at a concentration of 0.5mg/ml and 4mg/ml respectively. L20 was added to PduA* to give a 6:1 (peptide: hexamer) molar ratio. Circular dichroism was measured using a Chirascan^™^ spectrometer in a 0.05mm pathlength cuvette for all samples. CD spectra were analysed using the K2D3 software ^34^to estimate percentage helicity.

### Crystallography

PduL was crystallised using hanging drop vapour diffusion at a concentration of 8mg/ml. Crystals formed in 0.2M lithium sulphate, 0.1M sodium acetate, 50% PEG400 after approximately five days. Data were collected to 1.84Å at the Diamond light source I03 beamline using the PILATUS 6M detector. Data were processed using XIA2 and the space group P 3121 was assigned. Phases were calculated using molecular replacement with the model 5CUO, Phenix and Coot were used for all downstream processing and analysis^35,36^.

### Structural modelling

Models of a PduA* hexamer and pair of hexamers were produced using the structure 3NGK^9^, CABS-Dock was then run with either of these files as an input as well as either L20, P18 or D18. The simulation was run for 50 cycles and no preferred regions were selected to avoid any implicit bias. All structural figures presented were made using PyMOL^37^.

## Acknowledgements

This work was supported by the Biotechnology and Biological Sciences Research Council of the UK (BBSRC) strategic LoLa (BB/M002969/1) and an iCASE studentship from the BBSRC London Interdisciplinary Doctoral Programme (BB/T008709/1). We thank Simon Charnock and Prozomics for supporting the iCASE studentship. The authors would like to thank Diamond Light Source for beamtime and the staff of beamline I03 for assistance with data collection. We also thank Dr James Wright and Dr Ewan Main for donation of the vector pOPIN F(TEV).

