## Supplementary Information for "Enzyme encapsulation peptides bind to the groove between tessellating subunits of the bacterial microcompartment shell"

### Table of contents

**Figure S1**. Evidence from electron microscopy that the presence of encapsulation peptides or enzymes with encapsulation peptides prevents curvature of PduA sheets.

**Figure S2**. Circular dichroism spectra of the encapsulation peptides showing a helical structure is adopted when bound to PduA.

**Figure S3**. Conservation of the encapsulation peptide binding site in PduA, PduJ and EtuM.

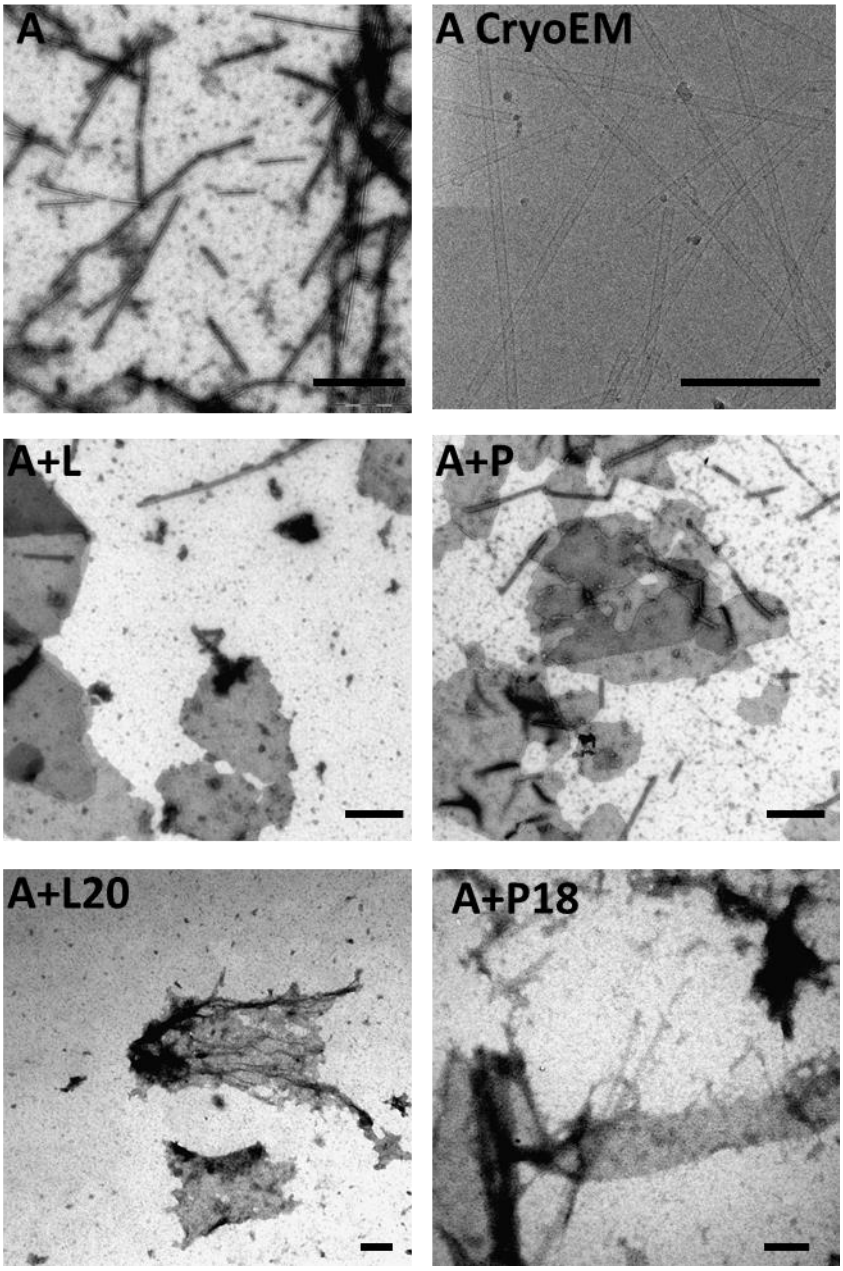


**Figure S1**. Evidence from electron microscopy that the presence of encapsulation peptides or enzymes with encapsulation peptides prevents curvature of PduA sheets. The top two panels show PduA alone in negative stain and in cryo conditions; the lower four panels show PduA in the presence of the enzymes PduL and PduP and their targeting peptides L20 and P18. PduA alone predominantly forms nanotubes, but in the presence of enzyme or peptide nanotubes are not favoured and sheets are dominant. This is consistent with encapsulation peptide binding at or near the hexamer-hexamer interface. The scale bar shown is 200nm.

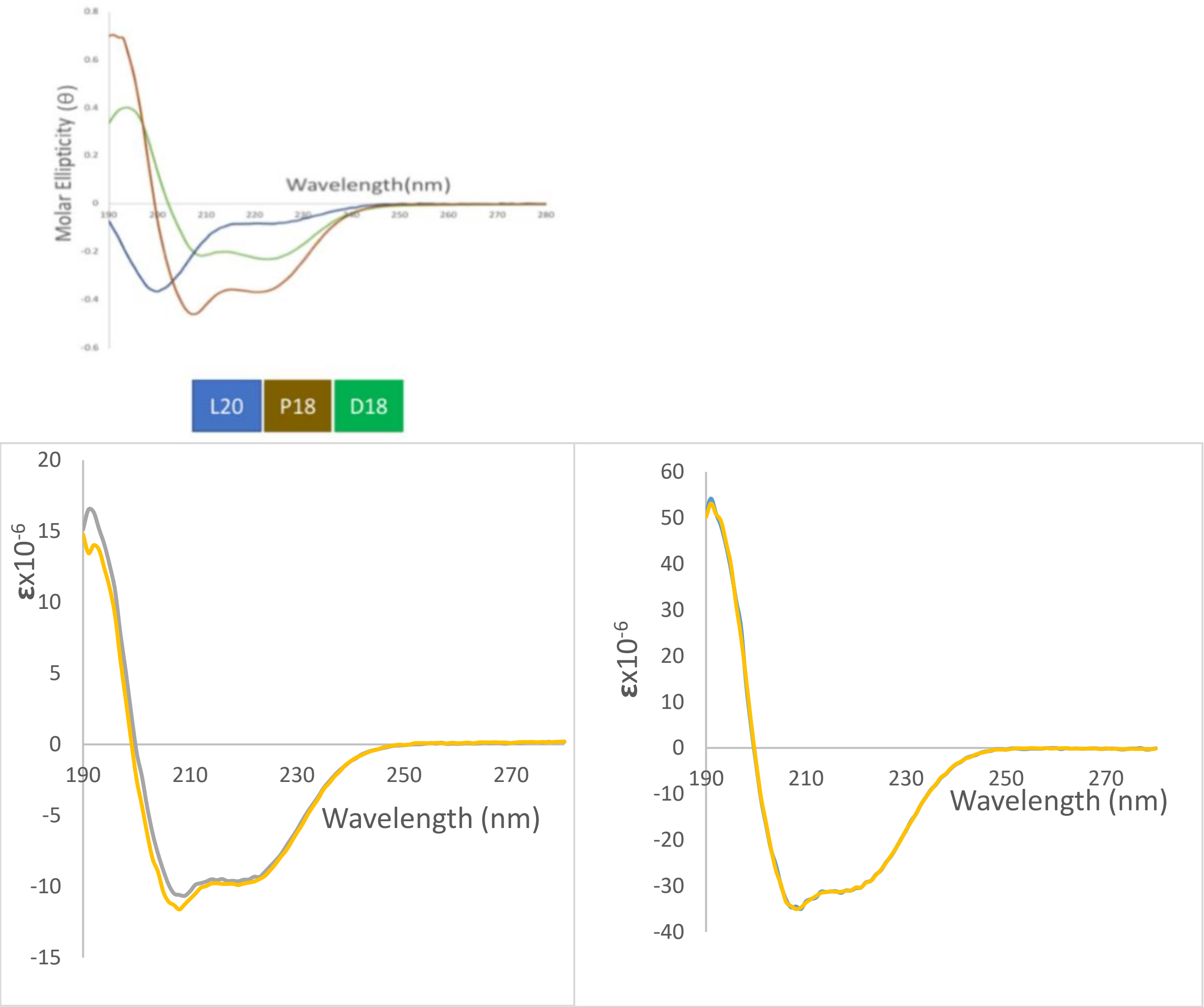


**Figure S2.** Encapsulation peptides are helical when bound to PduA. Assessed using circular dichroism spectroscopy. Peptides P18 and D18 are helical in solution compared to the largely unstructured L20 targeting peptide (top panel). The main spectra (left) show the increase in helicity when L20 binds to tessellating PduA (yellow) compared to the individual measurements of L20 peptide and PduA alone (grey), and the right show no change in helicity when PduL peptide is added to the non-tessellating variant PduA with GGSST linker (yellow both present, grey non-tessellating PduA only).

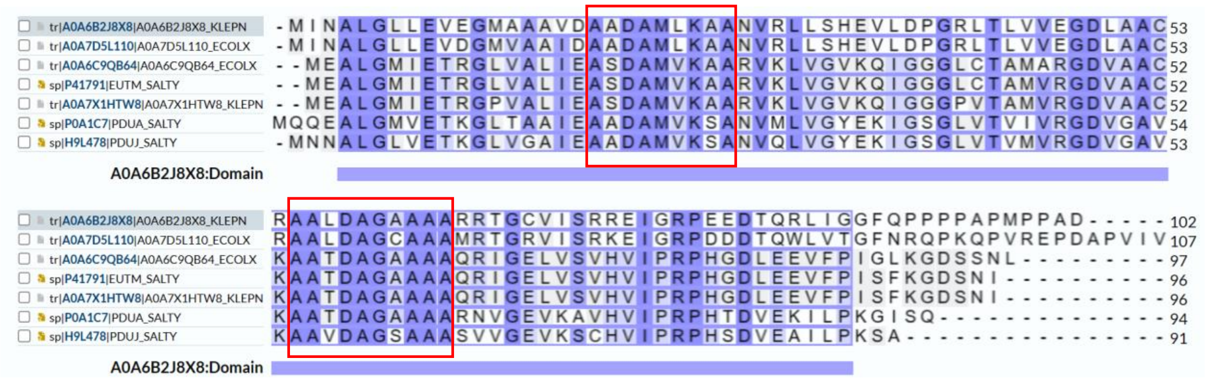


**Figure S3**. <u>Conservation of the encapsulation peptide binding site in PduA, PduJ and EtuM.</u> PduJ and PduA together account for over 70% of the shell proteins of the *Salmonella typhimurium* LT2 Pdu microcompartment. The conserved residues forming the encapsulation binding site on PduA are conserved in the major shell protein PduJ (see the conservation of binding site in the red boxes). The two signature sequences are on the surface of alpha-helices 1 and 2: AAD<u>A</u>$^{23}$MVK(<u>A/S)</u>$^{27}$A and <u>A</u>$^{56}$ATD<u>A</u>$^{60}$GA<u>A</u>$^{63}$AA, respectively (PduA, *Salmonella typhimurium* LT2, SALTY, numbering). The PduJ-PduJ interface will be most common and has the same hydrophobic groove as the PduA-PduA interface. Sequences shown in this figure are from *Salmonella typhimurium* LT2 (SALTY), *Klebsiella pneunomie* (KLEPN), and *E. coli* (ECOLX).
